# Data-driven trait heritability-based extraction of human facial phenotypes

**DOI:** 10.1101/2023.08.13.553129

**Authors:** Meng Yuan, Seppe Goovaerts, Hanne Hoskens, Stephen Richmond, Susan Walsh, Mark D. Shriver, John R. Shaffer, Mary L. Marazita, Seth M. Weinberg, Hilde Peeters, Peter Claes

## Abstract

A genome-wide association study (GWAS) of a complex, multi-dimensional morphological trait, such as the human face, typically relies on predefined and simplified phenotypic measurements, such as inter-landmark distances and angles. These measures are predominantly designed by human experts based on perceived biological or clinical knowledge. To avoid use handcrafted phenotypes (i.e., a priori expert-identified phenotypes), alternative automatically extracted phenotypic descriptors, such as features derived from dimension reduction techniques (e.g., principal component analysis), are employed. While the features generated by such computational algorithms capture the geometric variations of the biological shape, they are not necessarily genetically relevant. Therefore, genetically informed data-driven phenotyping is desirable. Here, we propose an approach where phenotyping is done through a data-driven optimization of trait heritability, defined as the degree of variation in a phenotypic trait in a population that is due to genetic variation. The resulting phenotyping process consists of two steps: 1) constructing a feature space that models shape variations using dimension reduction techniques, and 2) searching for directions in the feature space exhibiting high trait heritability using a genetic search algorithm (i.e., heuristic inspired by natural selection). We show that the phenotypes resulting from the proposed trait heritability-optimized training differ from those of principal components in the following aspects: 1) higher trait heritability, 2) higher SNP heritability, and 3) identification of the same number of independent genetic loci with a smaller number of effective traits. Our results demonstrate that data-driven trait heritability-based optimization enables the automatic extraction of genetically relevant phenotypes, as shown by their increased power in genome-wide association scans.

## I. Introduction

Understanding the genetic basis of human facial traits has important implications for various fields, including clinical genetics, forensic science, and evolutionary anthropology. Genome-wide association studies (GWASs) are a powerful tool to investigate the genetic architecture of complex traits such as facial features. A GWAS involves scanning numerous genetic variants across many individual genomes to find those statistically associated with a particular trait. While they have proven to be highly successful in identifying trait-associated genetic variants, the phenotypic descriptors in these studies are typically predetermined and often chosen because they are convenient or easy to obtain. Consequently, the effectiveness of GWAS in identifying true associations between traits and Single Nucleotide Polymorphisms (SNPs) can be influenced by the quality of the predefined phenotypic traits.

Facial shape phenotypes commonly used in GWAS include both traditional morphometric measurements [1]– [3] and more recent facial features acquired through computational methods [4], [5]. Traditional morphometric traits include direct measurements between key anatomical landmarks, such as distances, distance ratios, and angles. These measurements are often criticized for their limitations in capturing the full complexity of facial variation, as the multifaceted nature of facial shape is condensed into a constrained set of measurements. Furthermore, the definition and selection of key landmarks require biological or clinical knowledge and are typically determined by human experts. As alternatives, data-driven approaches, such as principal component analysis (PCA) and deep learning techniques of 3D facial surfaces, have gained popularity in facial phenotyping. These techniques enable the extraction of comprehensive measurements that can capture detailed facial surface topography without the need for predefining traits. While computational algorithms can generate features that capture the geometric variations of facial shapes, these features are not necessarily biologically meaningful or genetically relevant. Some studies [4], [5] have opted to retain a specific number of principal components (PCs) as phenotypes for GWAS, using the classical variance explained method. Other studies [6], [7] have assessed predefined facial features based on their heritability and chosen to retain those that demonstrate the highest heritability. Furthermore, a previous study [8] defined sibling-shared traits at an individual level, where a strong sibling resemblance also implies high heritability. To the best of our knowledge, no study has yet explored using trait heritability to actually optimize phenotypes at sample level, and to go beyond predefined features or measurements.

In this study, we present a data-driven, trait heritability-based phenotyping method that allows for the extraction of facial phenotypes in a genetically informed manner. The proposed method comprises two steps. Firstly, PCA is used to construct a lower-dimensional feature space that captures geometric facial shape variations. Subsequently, a genetic algorithm (GA; i.e., a heuristic inspired by natural selection) is employed as search strategy, focusing on trait heritability computed in a family-based design. The rationale behind this approach lies in the facial resemblance observed within families. Accordingly, it is assumed that genetically relevant variations are situated along the directions in the feature space where trait heritability is high. Thus, a GA is utilized to search for these directions by optimizing trait heritability. We trained our proposed phenotyping method using 3D facial surface scans of father-offspring pairs and then validated our approach on an independent dataset of parents-offspring trios. Additionally, we assessed the value of the derived facial traits in a GWAS of an independent cohort of unrelated European individuals with genome-wide SNP data. Our findings demonstrate that the phenotypes obtained using our trait heritability-optimization exhibit higher trait- and SNP-heritability compared to PCs in two independent cohorts. Furthermore, this new approach enables the identification of the same number of independent genetic loci with a smaller number of effective traits in GWAS. Therefore, the proposed method yields facial traits that contain a greater amount of genetically relevant information, thereby enhancing the efficiency of identifying genetic effects on facial shape in GWAS. In a broader context, these results offer an alternative perspective to non-heritability-optimized phenotyping, and our approach can be widely applied to any GWAS involving complex traits where access to phenotypic family data (e.g. parent-offspring, siblings, or twins) is available.

## II. Materials and Methods

### A. Data

Our study included two family-based cohorts and one population-based cohort.

1. The Avon Longitudinal Study of Parents and their Children (ALSPAC) [9], [10] is a UK-based family cohort study. The current study (B2409: “Exploring the heritability of facial features in fathers and offspring using spatially dense geometric morphometrics”) was approved by the ALSPAC Ethics and Law Committee and the Local Research Ethics Committees. Written informed consent was obtained from all children and their fathers. 3D facial images and self-reported information on demographic factors (e.g., sex, age, ethnicity), general physical characteristics (e.g., height, weight), and family relationship are available. Only individuals of self-reported European ancestry were retained for analysis. Participants with missing data on any of the covariates and those with poor-quality images were excluded. The resulting dataset consisted of 770 father-offspring pairs, which were used to train a GA.
2. The Technopolis dataset [8] is a Belgium-based family cohort. Ethical approval for the study was obtained from the Ethics Committee Research UZ/KU Leuven (S56392: ML10285). We applied the same quality control procedures as described for the ALSPAC data to maintain consistency. The final Technopolis study sample consisted of 163 parent-offspring trios, which served as an independent test set for evaluating the generalization ability of the trained GA model.
3. The EURO dataset [5] includes participants of European descent from two larger independent population-based cohort studies conducted in the United States (US) and United Kingdom (UK). Institutional review board approval was obtained at each recruitment site, and all participants gave their written informed consent before participation. Unlike the ALSPAC and Technopolis datasets, the EURO dataset consisted of unrelated individuals (n=8,246). In addition to 3D image data and relevant covariates, genotype data on individual SNPs were also available, allowing for the evaluation of extracted facial phenotypes through GWAS, for which this dataset was previously used [5]. This dataset only included participants without missing data and image artifacts. Information on the different genotyping platforms, imputation, and quality control can be found in [5]. Intersection of imputed and quality-controlled SNPs across the US and UK datasets yielded 7,417,619 SNPs for analysis.

### B. Pipeline

The proposed pipeline, shown in Fig 1, consists of two main steps: 1) dimensionality reduction using PCA, to establish a feature space that captures shape variations and extract compact representations from 3D facial images in contrast to using the original thousands of 3D surface vertices; and 2) a GA, to scan the feature space for directions exhibiting high trait heritability.

**Fig 1.**
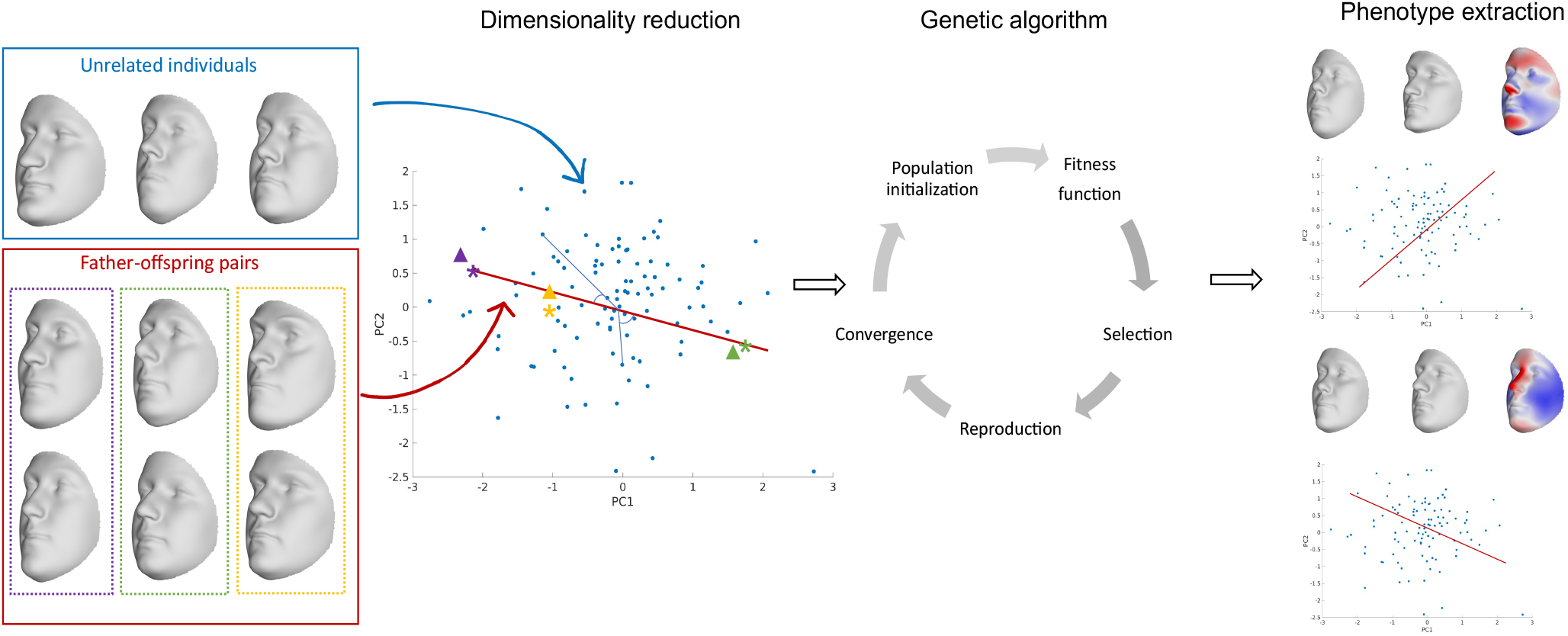
Pipeline illustrating the proposed approach for extracting human facial phenotypes based on trait heritability optimization, with 3D facial image as input and genetic relevant facial features as output. It consists of two steps: 1) constructing a feature space that models shape variations using dimension reduction techniques, and 2) searching for directions in the feature space where trait heritability is high using a genetic search algorithm.

#### 1) Dimensionality reduction

PCA is a standard linear technique for dimensionality reduction and has been widely used to model 3D facial shapes [11], [12]. To perform PCA, the 3D surface vertices are firstly transformed into a 2D matrix. Subsequently, a low-rank singular value decomposition (SVD) is performed on the mean-centered 2D matrix *X*, which is defined as *X* = *UΣV^T^*. The diagonal matrix *Σ* contains the singular values and the columns of U and *V* consist of the left and right singular vectors, respectively. The right singular vectors in *V* are the principal components. These components are orthogonal vectors, with the first component capturing the direction of maximum shape variance, while the subsequent components capture a decreasing proportion of variance. The later PCs with small variance carry little or noisy information about the data and can often be discarded without significant information loss. The optimal number of PCs is determined using parallel analysis [13]. Each face image is then represented as a combination of a reduced number of PCs, thereby preserving essential facial features.

#### 2) Genetic algorithm

Genetic algorithms are optimization methods inspired by the process of natural selection and evolution. Our objective was to search for *p* trait-heritability optimized directions in a *d* -dimensional PCA space (*p* ≤ *d*). To achieve this, we simulated *p* populations consisting of potential directions. For each population, we trained a GA model until the convergence condition was satisfied. The overall algorithm follows several steps (pseudocode in Table 1):

**Table 1.**
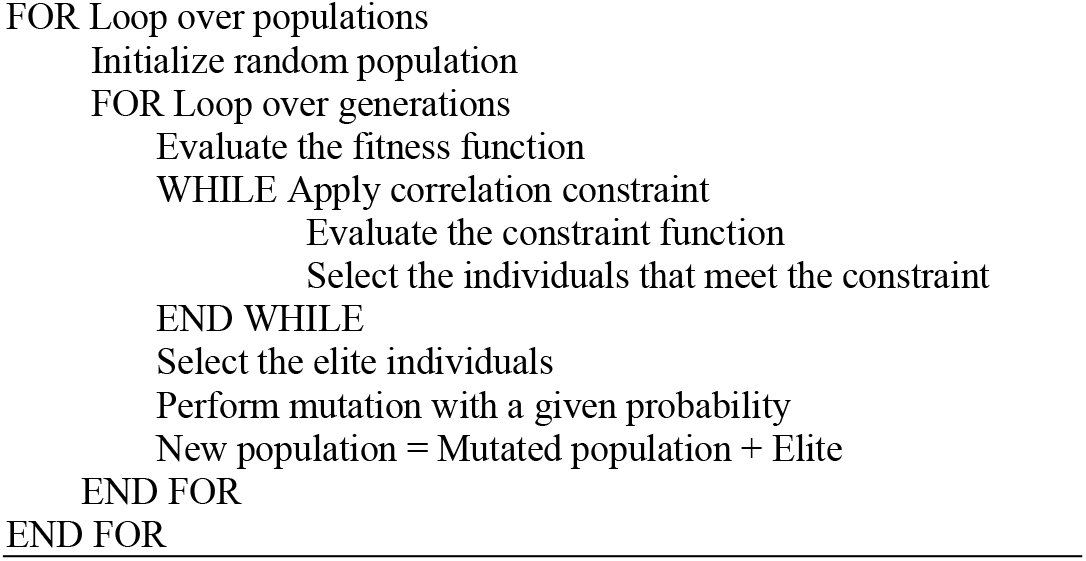
Genetic algorithm pseudocode

a. Initialization: The initial population 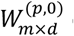 consists of *m* random directions in PCA space. Each random direction is a *d* -dimensional vector defined as a random linear combination of the PCs. Each direction corresponds to a distinct facial feature, and individuals can be scored along this direction. This is simply done by calculating the cosine angle between the individual’s vector and the vector of that direction in PCA space. Doing so, the cosine angle serves as a univariate trait score, quantifying the presence of a specific facial trait in an individual. Note that each PC on itself also defines a direction in PCA space, and therefore also codes for a facial trait, as previously done in facial GWAS [8]. Therefore, facial traits from individual PCs are used as a baseline to compare the optimized traits against.
b. Evaluation: Each direction is evaluated using a fitness function that quantifies its performance. The fitness function is defined using family-based trait heritability, indicating the degree to which the variability in a particular trait, as defined by the given direction, among individuals within a family can be attributed to genetic factors. This can be estimated by regressing offspring trait scores onto those of their respective parents [14]. Given the trait scores of parents and offspring, a linear regression (function regstats from Matlab 2022b) is performed to predict each univariate facial trait in children given the parents’ corresponding traits. The coefficient of determination (*R*^2^) reflects the trait heritability and is used to guide the evolution of the GA.
c. Selection: A subset of directions, i.e., elites, is selected based on their fitness values. Higher fitness implies a higher probability of selection.
d. Reproduction: To introduce diversity and explore new directions, all parent directions undergo a mutation operation to create new directions. The mutation operation introduces random changes to directions. In the *g* -th generation, the new population 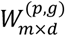 consists of the elite directions and mutated directions.
e. Constraint: When the GA model incorporates the correlation constraint (GAC), it selectively retains a subset of the current directions that exhibit mean correlations with the previous best directions *B*(*p*−1)×*d* that are less than or equal to a predetermined threshold 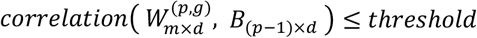. Such a constraint is not standardly used in GA and is introduced here to generate subsequent optimized directions that show a level of distinctiveness from other already optimized directions. The assumption is that without such a constraint, multiple rounds of the GA would result in the same and therefore redundant directions.
f. Termination: The algorithm proceeds with new generations until it reaches a predefined maximum number of generations.

Through iterative application of the described steps, the GA systematically explores the feature space, gradually evolving the directions with a tendency towards increased trait heritability.

### C. Evaluation metrics

#### 1) Trait- and SNP-Heritability

In addition to assessing the generalization ability of the trained GA model by evaluating trait heritability usin*g* an additional family-based dataset (i.e., Technopolis), we also investigated whether phenotypes derived from the trait heritability-optimized method yield genetically informative traits for GWAS. Therefore, we compared the SNP-based heritability of different facial traits as deducted from GWAS using linkage disequilibrium score regression (LDSC) (published software https://github.com/bulik/ldsc/) [15]. SNP-heritability is defined as the proportion of phenotypic variance that is explained by additive genetic effects of SNPs. First, SNPs were intersected with the HapMap3 [16] SNPs and any SNP with non-matching alleles was removed, as well as SNPs within the major histocompatibility complex region. The SNP heritability of each univariate trait was then estimated by LDSC using the GWAS summary statistics of the EURO dataset. European-derived LD scores were used in LDSC (downloaded from https://doi.org/10.5281/zenodo.7768714).

#### 2) Identification of trait-associated genetic loci

Aside from using GWAS outcomes in LDSC, we also investigated the number of genetic loci identified in a single or a group of GWAS by peak detection. Given a group of univariate traits (e.g. PCs or multiple optimized traits) and their respective GWAS summary statistics, for each SNP the minimal P-value was first retained. For a single trait, the typical peak detection threshold in GWAS is the genome-wide significance threshold (P < 5 x 10^−8^). However, for a group of traits, adjustment for the multiple testing burden is needed and this was done by dividing the genome-wide threshold by the number of independent univariate phenotypes (i.e., effective traits). We estimated the number of independent phenotypes using permutation testing [17]. Specifically, each of 7,417,619 SNPs was randomly permuted and the same GWASs were repeated. This allowed an estimate of the null-distribution of the minimum P-values for each SNP across the univariate traits. The number of independent phenotypes was then estimated as 0.05 divided by the 5th percentile of this null distribution. Subsequently, peak calling was performed in three steps, starting with the SNPs that reached the adjusted genome-wide significance threshold. First, all SNPs within 250 kb of the most significant SNP, as well as those within 1 Mb and in LD (*r*^2^ > 10^−2^) were clumped into a single locus represented by the most significant (lead) SNP. This was repeated until all SNPs were assigned to a locus. Next, any two loci were merged if the representative lead SNPs were within 10 Mb and in LD (*r*^2^ > 10^−2^). This locus was then represented by the SNP with the lowest P-value. Lastly, any peaks represented by a single SNP below the adjusted genome-wide significance threshold were disregarded to improve robustness.

#### 3) Phenotypic variation explained by facial traits

For a single or a group of facial traits selected or optimized as directions in PCA-space, we measured the amount of phenotypic variation that they capture from the original 3D facial images. We employed a partial least-squares (PLS) regression (function plsregress from Matlab 2022b) to predict the original images or 3D surface vertices based on these traits. The sum of variances of all PLS components reflects the phenotypic variance explained.

### D. Implementation details

#### 1) 3D facial image processing

First, we adhered to the procedures outlined previously [5] to perform spatially dense facial quasi-landmarking and quality control using the MeshMonk toolbox [18]. Subsequently, the quality-controlled images, comprising 7,160 dense quasi-landmarks were aligned using generalized Procrustes analysis (GPA), symmetrized, and were further adjusted for potential confounding factors such as sex, age, age squared, height, weight, and facial size in a PLS regression. Both the ALSPAC and EURO datasets underwent further correction for the camera system used. Moreover, for the EURO datasets, an additional correction for population structure was implemented by including the first 4 genomic PCA axes (i.e., ancestry axes) in the PLS regression.

#### 2) Model training

The first step involved reducing the dimensionality of the original surface meshes, which consist of 7,160 dense quasi-landmarks, and constructing a lower-dimensional feature space. In order to capture a substantial amount of facial variation, we applied PCA on the facial images from the EURO dataset, which offered a large sample size. Subsequently, the facial images from the ALSPAC and Technopolis datasets were projected onto this feature space. To determine the number of PCs needed to adequately summarize shape variation, we used parallel analysis [13], resulting in 70 PCs which were then normalized to unit variance.

In the second step, we applied a GA to search for directions in the feature space that exhibit high trait heritability. The initial population consisted of 1,000 randomly generated directions within the feature space. In each generation, the top 5% of the total population, i.e., elites, were selected based on their fitness values and retained for the next generation of evolution. All parent directions underwent a mutation operation to create new directions where Gaussian noise within a range of 10^−4^ of variance was added. For the regular GA model, a total of 70 populations were created, with each population evolving independently for over 500 generations. In contrast, for the constrained model (GAC), the training process was conducted sequentially, and the best directions from previously trained populations were saved. Only a subset of the current directions with mean correlations equal to or less than 0.1 with the previous best directions were retained during evolution. After each generation, the correlation threshold was relaxed by 10^−4^ for the subsequent generation, preventing the algorithm from becoming overly constrained.

#### 3) Genome-wide association meta-analysis

For each univariate trait, GWASs were conducted in the US and UK cohorts independently using linear regression (function regstats from Matlab 2022b) where SNPs were coded under the additive genetic model (0, 1, 2). This yielded effect size and standard error estimates for the US and UK cohort independently, which were then meta-analyzed using the inverse-variance weighted method [19]. Meta P-values were obtained using a two-tailed test.

In order to validate and compare the added value of our proposed phenotyping protocol we investigated four trait groups. The first group consists of PCs (PCA), the second group of traits are random directions in PCA space (RANDOM), the third group of traits are directions optimized in trait-heritability using the proposed genetic algorithm (GA), and finally the fourth group of traits comprises directions optimized in trait-heritability with the constraint of trait diversity added in a subsequential optimization scheme (GAC). To investigate the number of identified genetic loci under different numbers of traits, we gradually increased the absolute numbers of traits in each trait group. The experiments were conducted with absolute numbers of traits equal to 1, 6, 10, 20, 30, 40, 50, 60, and 70. We then combined multiple GWASs on a group of univariate traits and detected significant peaks. For PCA and GAC, the derived traits are arranged in a specific order, i.e., descending explained variance and an order decided via the correlation constraints in subsequent rounds of optimization, respectively. However, the GA traits and traits along randomly selected directions are not presented in any particular order. Therefore, for these two groups, the results of multiple randomly sampled sets of traits were generated. Specifically, we randomly sampled a particular number of traits, identified independent peaks based on these traits, and repeated this process 50 times. Subsequently, the median and standard deviation of the results were represented as error bars.

## III. Results

We evaluated the genetic relevance of the facial phenotypes obtained through our trait heritability-optimized approach in the following aspects: 1) trait heritability, 2) SNP-based heritability, and 3) the effectiveness of identifying independent genetic loci associated with the facial phenotypes through GWAS.

### A. Heritability

In Fig 2.A-B, it can be observed that for both the training data (ALSPAC) and testing data (Technopolis), the traits extracted by GA and GAC exhibited higher median trait heritability, compared to PCs and traits along randomly selected directions. GA generated the highest median heritability followed by GAC. There were no statistically significant differences in median heritability between PCs and traits along randomly selected directions. In addition, in Fig 2.C, the traits extracted by GA and GAC consistently demonstrated higher median SNP heritability in comparison to all other methods. GA generated the highest median SNP-heritability followed by GAC. However, as seen in Fig 3, the GA traits exhibited substantial overlap in facial regions (e.g., tip of the nose and nostrils), indicating a lack of diversity. Consequently, the recurring occurrence of these highly heritable traits by GA results in an inflation of the median heritability, contrasting with the constrained outcomes obtained from GAC.

**Fig 2.**
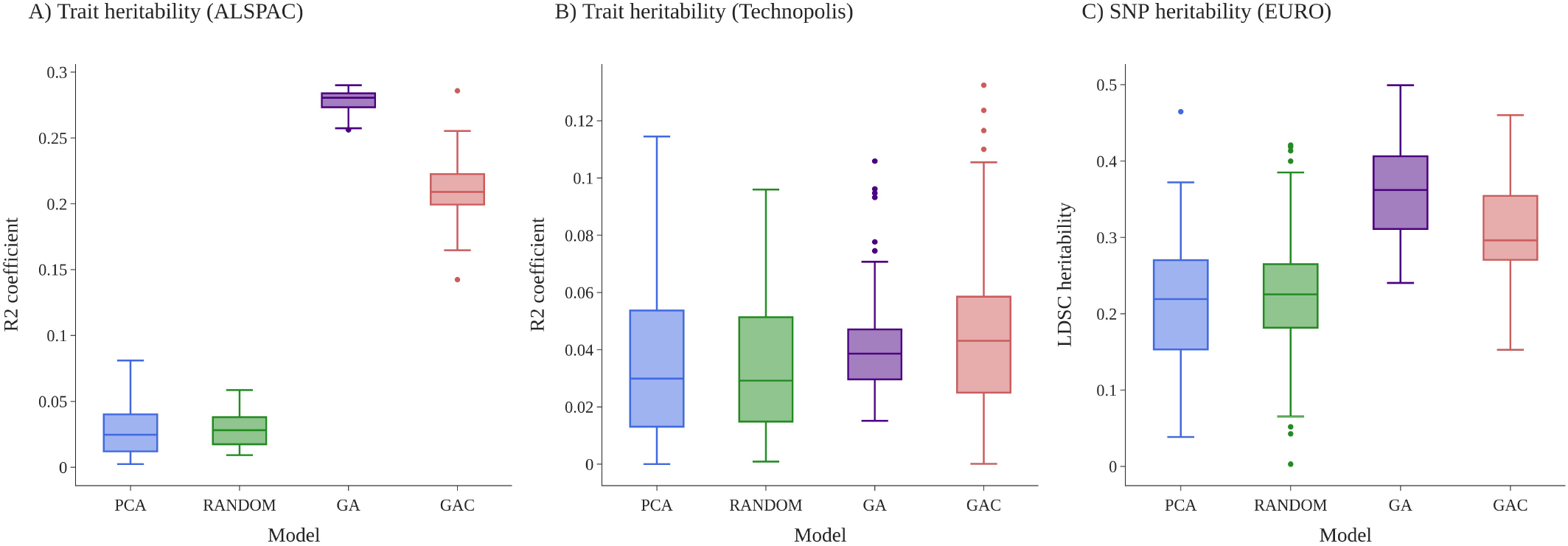
Comparing facial traits in terms of heritability: Trait heritability for (A) ALSPAC; training data, (B) Technopolis; testing data, and (C) SNP-based heritability estimated from GWAS results obtained from a cohort of unrelated individuals; EURO. The colors represent the phenotypes, blue for PCs, green for traits along randomly selected directions in the feature space (RANDOM), purple for traits extracted by GA, and red for traits extracted by GAC.

**Fig 3.**
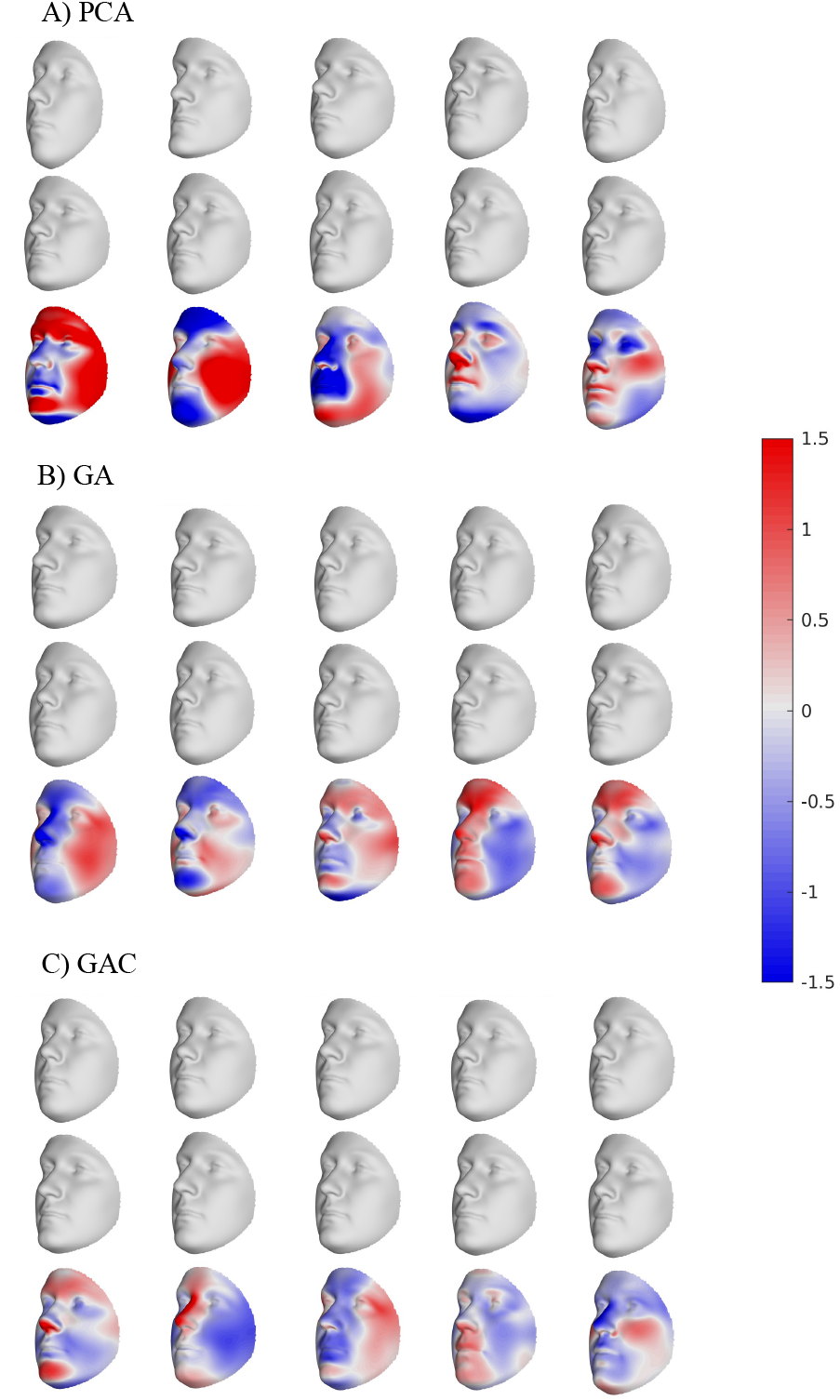
Visualization of facial phenotypes extracted by (A) PCA, displaying the first 5 principal components with variations explained in decreasing order from left to right, (B) GA, and (C) GAC, showing selected traits with distinct characteristics. The shown shape variation ranges from μ-3*σ (upper) to μ+ 3*σ (middle), where μ and σ are the mean and standard deviation along the dimension of interest, respectively. The lower row displays a colormap showing the difference between the first two rows expressed in mm.

### B. Trait-associated genetic loci

Fig 4.A illustrates the completeness or distinctiveness of the facial traits in a group by presenting both the absolute number of traits (i.e., those submitted to GWASs) and the relative number of traits (i.e., the number of independent traits calculated by permutation test). The traits extracted by GA exhibited a high degree of correlation, resulting in only 14 independent dimensions from 70 dimensions. On the other hand, the traits extracted by GAC showed relatively lower correlation, yielding 39 independent dimensions from 70 dimensions. In contrast, traits along randomly selected directions and PCs showed the mostly uncorrelated patterns. In other words, they better span the complete scope of facial variation, by not just focussing on genetically interesting traits and e.g., also describing non-genetic face differences.

**Fig 4.**
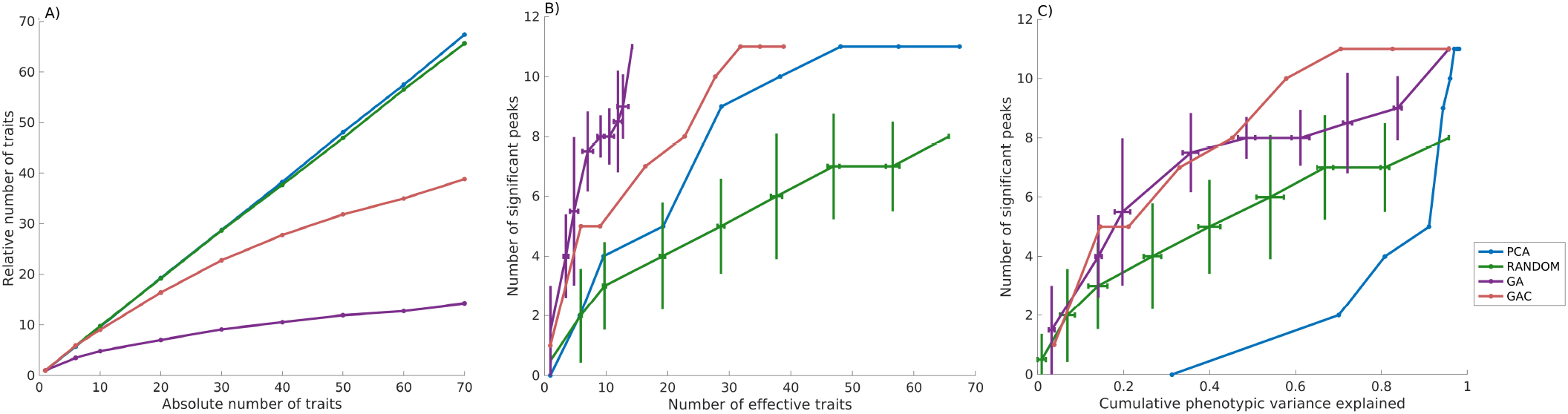
Comparing facial phenotypes in terms of (A) the number of effective traits, (B) the effectiveness of identifying independent genetic loci through GWAS, and (C) the phenotypic variation captured by traits and their corresponding number of significant genetic loci in GWAS. The colors represent the phenotypes, blue for PCs, green for traits along randomly selected directions in the feature space (RANDOM), purple for traits extracted by GA, and red for traits extracted by GAC.

Fig 4.B presents the number of independent genetic loci identified when aggregating multiple univariate GWASs, alongside the number of effective traits. As expected, the inclusion of more effective traits resulted in an increase in the number of independent genetic loci across all methods. In comparing curves, the better behavior involves a steeper curve indicating that for a type of traits a lesser number of effective traits can result in more genetic loci. The curve representing the cumulative number of peaks based on GA traits exhibited the steepest slope, reaching the maximum number of significant peaks (n=11) when 14 effective traits were included. Following that were the curves based on GAC and PCA, which reached the same maximum number of significant peaks with 39 and 67 effective traits, respectively. The curve resulting from traits along randomly selected directions situated lowest, identifying the smallest number of genetic loci (n=8). It is noteworthy that the results of traits along randomly selected directions and GA traits exhibited substantial variation due to the different random selections.

Fig 4.C demonstrates the relationship between the phenotypic variation captured by the types of traits and their corresponding number of significant genetic loci in GWAS. The patterns observed in GA and GAC exhibited similarities, with the fastest increase of the number of identified genetic loci occurring when the absolute number of traits was smaller than 30. This indicates that although the resulting traits may capture fewer geometric facial variations, they yield a greater discovery in GWAS. On the other hand, the first PC explained 31.22% of the phenotypic variation but led to zero significant genetic loci. The identification of two independent genetic loci only became possible when considering the first 6 PCs, which in total captured 70.05% of facial variations. The curve of PCA exhibited a rapid increase starting from 6 PCs and reaches its maximum number of peaks at 50 PCs. Interestingly, this suggests that while a substantial amount of geometric phenotypic variance is captured by the first set of PCs, it may not necessarily correspond to genetically relevant information. This observation is further supported by the visualization of facial features presented in Fig 3. Specifically, the facial phenotypes represented by the first 5 PCs encompassed a large region of the face. In contrast, the GAC traits particularly targeted smaller, known to be heritable facial regions, such as the nasal and zygoma regions. Furthermore, the curve of traits along randomly selected directions situated between those of GA-optimized approaches and PCA, increasing gradually and reaching the smallest number of significant peaks among all methods.

## IV. Discussion

In this work, we proposed a framework for facial phenotyping based on a trait heritability-optimized approach to identify genetic variations influencing facial shape. The proposed methodology involves constructing a low-dimensional feature space of 3D facial scans using PCA, followed by applying a genetic algorithm to search for directions within this feature space exhibiting high trait heritability. As anticipated, the phenotypes obtained through the trait heritability-optimized training exhibited higher trait heritability and SNP heritability compared to principal components. Moreover, this new approach facilitates the identification of an equivalent number of independent genetic loci with fewer effective traits in GWAS as compared to PCA or random traits. In summary, the proposed data-driven, trait heritability-optimized method, provides an alternative to the prevailing hypothesis-driven subjective selection of phenotypes, and yields highly efficient and genetically informative facial traits for GWAS analysis.

Although PCA is a powerful tool for reducing the dimensionality of datasets with high-dimensional correlated information, the genetic value (e.g., discovery rate in GWAS) or more general the biological value of its extracted features is questionable [20]. There were no statistically significant differences in median heritability between PCs and traits along randomly selected directions in the PCA space (Fig 2). Furthermore, compared to the first few PCs, traits along randomly selected directions captured a smaller proportion of shape variance but identified a larger number of significant genetic loci (Fig 4.C). These observations indicate that the geometric variations in facial shape captured by PCs are not strongly indicative of underlying genetic factors. This can be attributed to the fundamental characteristic of PCs, which are mathematical constructs designed to capture as much shape variation as possible. Each PC is inherently bound by a specific direction of variation, thereby ensuring that the PCs are orthogonal to one another and arranged in a descending order of shape variance. In contrast, traits along randomly selected directions or GA-optimized traits have the freedom to be correlated and demonstrate substantial associations with genotypes simultaneously.

The idea of selecting heritable traits using phenotype-only designs (e.g., parent-offspring, twins, etc.) has been around for a while, even from before the genomic-era. One common point of discussion in phenotype-based heritability studies, is the inability to transfer heritability estimates from one population sample to another [21]. Therefore, even though the idea of using phenotypes with a high trait-heritability score in subsequent genetic studies has been around for decades, they never yielded a convincing added value over other phenotypes. Common to all studies to date, is that phenotypes remain pre-selected, and that trait-heritability is only used subsequently to prioritize or rank traits. In contrast, in this work, trait heritability is used from the start, and drives the selection of phenotypes. We believe it is this optimization, instead of simple ranking, that makes the difference, and allows for a better transferability of trait scores from one population sample to another. This is illustrated in our results, using two independent cohorts in which the optimized traits remain heritable.

The concept of scoring traits on a specific direction within the PCA space was utilized in a previous study [8] where researchers conducted GWAS based on sibling-shared traits. Inspired by this methodology, we first investigated whether traits along randomly chosen directions in the PCA space could serve as alternative traits to principal components. While traits along randomly selected directions demonstrated comparable median heritability, they resulted in a lower number of significant genetic loci in GWAS compared to PCs, when taking all PCs into account (Fig 4.B). However, traits along randomly selected directions capturing less shape variance only identified more genetic loci compared to the first few PCs (Fig 4.C). Building upon these findings, we took a step further by implementing a trait heritability-optimized approach, which was initialized by a set of randomly selected directions. We then applied GA to systematically scan all possible directions within the PCA space, aiming to search for genetically relevant directions. As hypothesized, this approach led to a reorganization of the feature space, emphasizing traits with phenotypic distributions that were most consistent with underlying genetic factors. The results demonstrated that, when the number of dimensions was limited, the GA-optimized traits yielded a higher number of significant genetic loci compared to those obtained through PCs. Moreover, as the number of effective traits increased, PCA and GA-optimized methods reached the same saturation point in terms of the number of identified peaks. This is because we employed a two-step approach, where GA-optimized approaches were constructed based on a pre-determined PCA feature space. Thus, it would be interesting for future studies to investigate an end-to-end approach that integrates dimension reduction and trait-heritability optimization simultaneously.

We also proposed a constrained version of the GA model (GAC), alongside the regular GA model. The motivation behind this exploration was that the traits generated by the standard GA exhibited limited diversity. I.e., the same and similar traits were extracted, time and again. In contrast, the incorporation of correlation constraints within the GAC model resulted in more diverse facial phenotypes as seen in Fig 3. Therefore, adding the constraints allows for capturing more of the geometric facial variation as seen in Fig 4.C. Therefore, the GAC model combines the heritable traits using GA, with a range of diversity as seen in PCs.

In conclusion, our proposed data-driven, trait heritability-optimized approach offers an opportunity to automatically extract genetically relevant phenotypes, as shown by their increased power in GWAS. This methodology can be seamlessly applied to the study of other types of images, such as the brain or the skull. Furthermore, the algorithm presented in this study can be extended to other applications by modifying the fitness function of the genetic algorithm. For instance, it can be adapted to extract quantitative features specifically related to rare genetic variants. This flexibility allows for the potential applications of our approach in a wide range of genetic studies.

## ACKNOWLEDGMENT

We are extremely grateful to all the families who took part in this study, the midwives for their help in recruiting them, and the whole ALSPAC team, which includes interviewers, computer and laboratory technicians, clerical workers, research scientists, volunteers, managers, receptionists and nurses. We are also very grateful to all of the Technopolis and EURO participants and lab members for generously donating their time, and Technopolis Belgium for allowing the use of their facilities for this research.

## Contributions

M.Y. and S.G. under supervision of P.C. developed the methodology and designed the experiments. M.Y. performed the computations and drafted the manuscript. All authors discussed the results and contributed to the final manuscript.

